# Flow arrest in the plasma membrane

**DOI:** 10.1101/575456

**Authors:** Michael Chein, Eran Perlson, Yael Roichman

## Abstract

The arrangement of receptors in the plasma membrane strongly affects the ability of a cell to sense its environment both in terms of sensitivity and in terms of spatial resolution. The spatial and temporal arrangement of the receptors is affected in turn by the mechanical properties and the structure of the cell membrane. Here we focus on characterizing the flow of the membrane in response to the motion of a protein embedded in it. We do so by measuring the correlated diffusion of extracellularly tagged transmembrane neurotrophin receptors TrkB and p75 on transfected neuronal cells. In accord with previous reports, we find that the motion of single receptors exhibits transient confinement to sub-micron domains. We confirm predictions based on hydrodynamics of fluid membranes, finding long-range correlations in the motion of the receptors in the plasma membrane. However, we discover that these correlations do not persist for long ranges, as predicted, but decay exponentially, with a typical decay length on the scale of the average confining domain size.

## Main

The plasma membrane is a highly dynamic heterogeneous object. It is made of a lipid bilayer, in which a variety of proteins are embedded, and is anchored to the cytoskeleton through binding domains ^1-6^. Although the weight percent of lipids and proteins in the plasma membrane is comparable, the conventional description of the plasma membrane is of a complex fluid, in which protein inclusions can diffuse^7^. This point of view is supported by the observed mobility of entities of different sizes embedded in the plasma membrane, such as protein aggregates and transient lipid rafts^6-10^. For this reason, models such as the Saffman-Delbruck model^11^ that describe the plasma membrane as a simple viscous fluid, are often inconsistent with experimental observations. For example, a significantly lower mobility of proteins is measured in live cells as compared to in artificial lipid-based vesicles that are composed of a fluid lipid bilayer^3,8^ and in red blood cells^12^. The question then arises whether a membrane that is “more mosaic than fluid” ^3^ can flow like a fluid lipid bilayer. Continuous flow of the plasma membrane is believed to be the mechanism for fast equilibration of membrane tension. Such flow is therefore essential for the commonly suggested mediation of fast long-range mechanical signals within cells (see for example and reference in). Surprisingly, a recent study by Shi *et al.* has put forth proof that local changes in the cell membrane tension result in localized mechanical signaling. In these experiments, short tethers where pulled out of cells to perturb locally the tension in the cell membrane and to measure it. No mechanical coupling between two tethers drawn at distances of 5-15 nm was observed in contrast to the strong coupling found for tethers drawn from cell attached membrane blebs. To account for their findings, the authors suggest a model in which the membrane has a gel like structure, and that trans-membrane proteins anchored to the cytoskeleton are the source of the membrane resistance to flow.

The correlated motion of tracer particle gives a direct measure of fluid flow. It was proposed and used to characterize the viscoelastic properties of complex fluids ^22-24^ in microscopic length scales. In this method, the effect of the mechanical perturbation created by the thermal motion of one tracer on the motion of a second tracer is measured as a function of their separation. In order for correlations between the motion of the two tracers in a membrane to exist, information through flow or deformation of the membrane needs to pass between them ^24-28^. As a result, the correlated motion is not only a means to characterize the viscoelastic properties of a complex fluid at large length scales, that are larger than the scale of structural and compositional inhomogeneity in it, but also a way to characterize the relevant length scale of heterogeneity^23,29,30^. This measurement is insensitive to tracer size or to tracer confinement, as long as the material in which the tracer is suspended can flow or deform ^24,31^.

Here we use single particle tracking (SPT) of trans-membrane proteins diffusing on the plasma membrane of two types of cells to extract the characteristic flow of the plasma membrane. From this direct measurement we quantify the characteristic flow of the plasma membrane and its range before being arrested. Our results further support and provide new insight into the surprising results of Shi *et al.*^21^.

Our experiments consisted of imaging the motion of TrkB and p75 receptors along the plasma membrane of neuronal and Hek 293T cells. The receptors were tagged extracellularly using ACP-CoA surface labeling, previously described in^39^, (Figure 1 **A,B)** resulting in minimal background noise arising from the cell cytoplasm. Importantly, the tagged receptors keep their biological function (see supplemental materials and^39^). We use TIRF to image the motion of these receptors on the cell membrane on a single molecule level (Figure 1**C)** and extract their motion using video microscopy ^40^. To obtain significant statistics we analyze over 40,000 trajectories each of them longer than 20 frames, i.e. at least 0.5 s long, from at least 40 different cells, from several independent cultures for each receptor and cell type.

**Figure 1:**
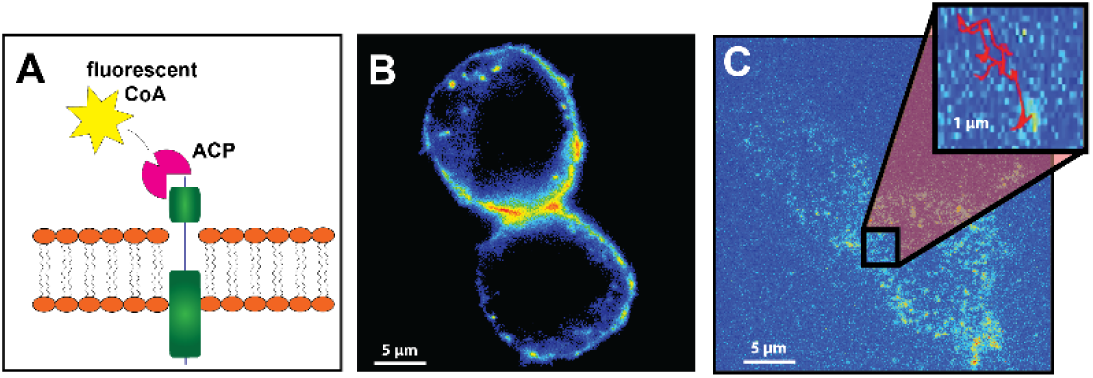
Experimental system. **A** A schematic representation of the TrkB receptor and its fluorescent tagging. **B** Epifluorescence image of infected primary motor neuron showing that TrkB ACP is localized on the cell membrane. **C** Single TrkB receptors on the cell membrane of a motor neuron as imaged by TIRF.

From the extracted trajectory of the receptors we calculate the correlated displacement of two proteins *α* and *β* according to:

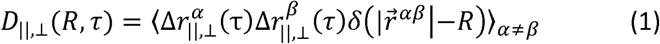

where 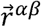 is the vector connecting the two proteins, 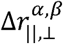 is the displacement of protein *α, β* along the direction parallel or perpendicular to 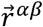. The correlated displacement of both receptors, TrkB and p75 in both cell types with *τ* = 25 *ms* is presented in Figure 2.

**Figure 2:**
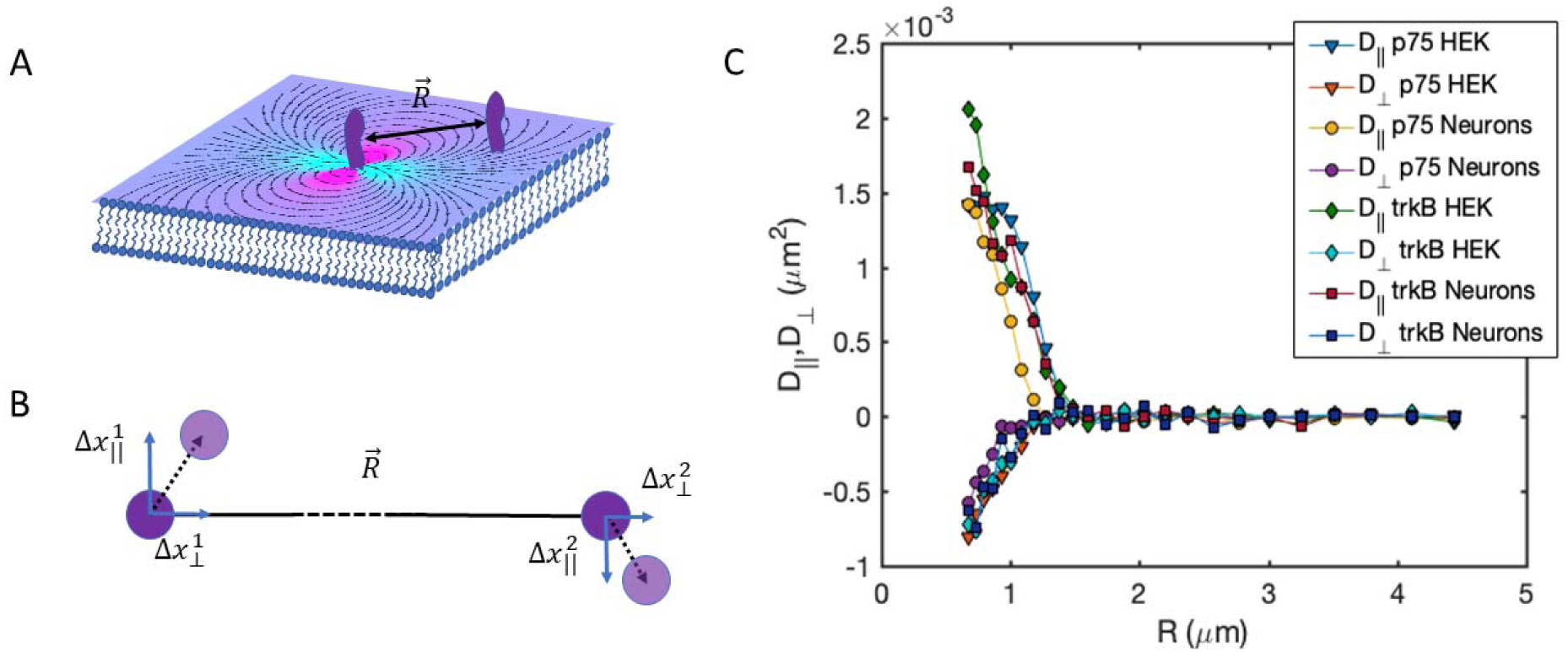
Two point microrheology of trans-membrane proteins in the plasma membrane. **A)** Illustration of two proteins embedded in the plasma membrane at a distance 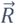 **B)** Cross correlation between displacements along the line connecting the proteins and the line perpendicular to them are calculated. **C)** the measure correlated diffusion of P75 and TrkB in Neurons and in HEK cells as a function of distance. All the results fall on the same lines confirming the similar flow characteristics of the plasma membrane in response to the different proteins and in the different cell types.

In a membrane with mobile protein inclusions suspended in fluid, for protein separations larger than the protein diameter, *r* ≫ *a,* we expect 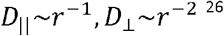. However, if the membrane is supported on a glass surface, which is the case in total internal reflection fluorescence microscopy (TIRF), for large separations, we expect 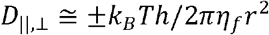, where *k*_*B*_*T* is the thermal energy, *h* is the distance between the membrane and the supporting surface, and *η*_*f*_ is the viscosity of the surrounding fluid. Similarly, immobile inclusions in the plasma membrane will cause the correlated diffusion in both directions to decay at large separations as *r*^−2 37^.

As expected, the correlated diffusion in the parallel and transverse direction of both receptors and in the different cell types agree well (Figure 2C). In order to compare these measurement to the theoretical prediction of Oppenheimer and Diamant^38^ we estimated the parameters entering the calculation as follows: the distance of the plasma membrane to the glass slide to be *h* = 50 nm ^44,45^, the thickness of the plasma membrane to be w = 8 nm ^4S^, the radius of the trans membrane part of the receptors to be *a*= 2 nm, and the viscosity of the cytosol and surrounding medium to be equal and *η*_*F*_ = 0.001 Pa.s (similar to water). In addition, we extracted the self-diffusion of the receptors from single particle data and obtained *D* = 0.375 *μm*^*2*^/s, from which we calculated the viscosity of membrane according to the Stokes Einstein equations to be *η*_*m*_ = 0.291 Pa.s.

The calculated correlated diffusion agrees well with the experimentally measured one for both receptors, with positive longitudinal correlations and negative transverse correlation (Figure 3). Surprisingly, these correlations seem to decay much faster than expected. In order to account for this discrepancy, we modify the theoretical prediction by multiplying the transverse and longitudinal solutions with the same exponentially decaying function. We then extract from the fit a typical decay length of 450 ± 80 nm (Figure 3 **A** and **B).** This implies that the flow of the plasma membrane is completely arrested on length scales larger than 1.5 ± 0.2 nm, as see in Figure 3. We note that the fact that the membrane flow is arrested at large distances does not imply that proteins and other biomolecules cannot diffuse in between domains. These findings indicate that the motion of proteins in the plasma membrane is strongly correlated at intermediate distances (approximately 450 nm) via the flow field induced in the membrane by their motion. However, at larger distances (approximately 1.5 nm and larger) the motion of proteins is completely decoupled.

**Figure 3:**
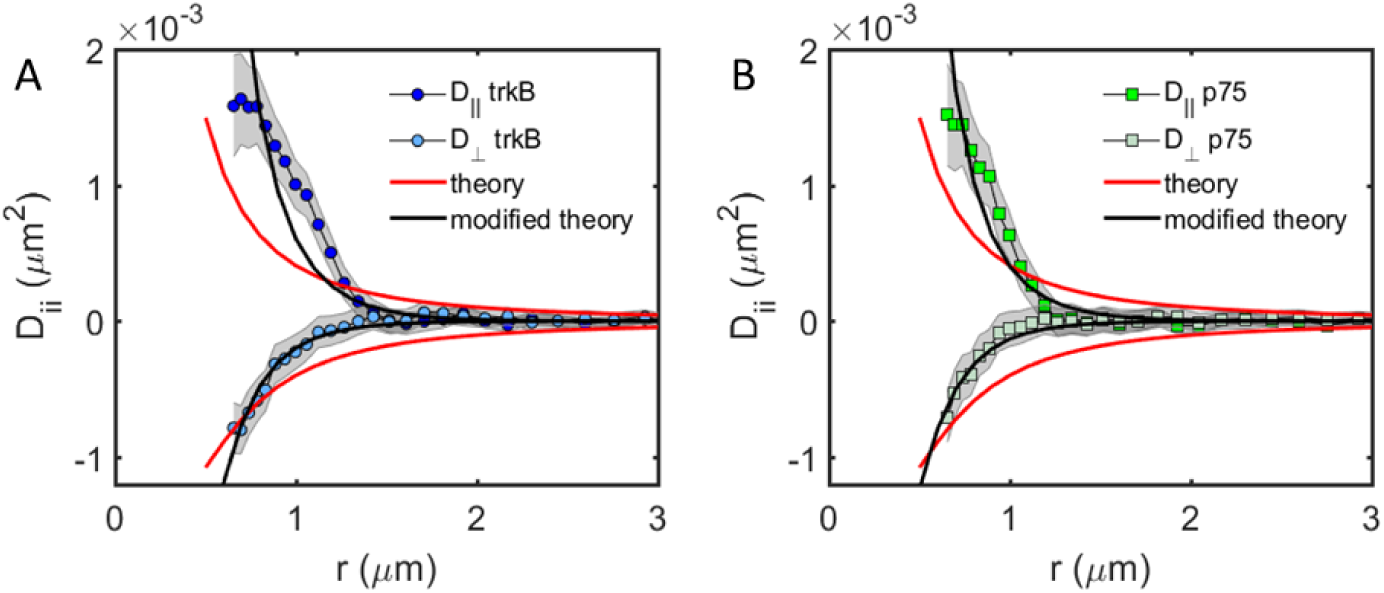
Correlated diffusion of protein receptors on the plasma membrane of neuronal cells exhibit long-range correlations. Comparison between the correlated diffusion of TrkB (**A**) and p75 (**B**) in the longitudinal and transverse directions. Similar behavior is observed for both receptors with a qualitative agreement with the simplified theory of^38^ (red solid line, see text for the parameter choice for the theoretical curves). The experimental results of both proteins fit better the modified theory (black solid line)

To relate the decay length of correlation in protein motion to structural features in the membrane we analyze the single protein stochastic motion of the proteins. Typical trajectories of receptors in the plasma membrane exhibit an alternation between two modes of motion, confined and free (Figure 4 **A,B).** The transient confinement events are similarly manifested in the MSD of both TrkB and p75 receptors (Figure 4C). The ensemble averaged MSD, ⟨Δ*r*^*2*^⟩_*En*_, of 3 s long trajectories and the MSD curves of both receptors agree well with each other. The MSD of a protein diffusing in a bounded domain is given by 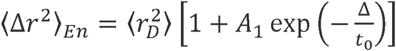, where 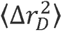 is the average domain area, *A*_1_is a geometrical factor, and *t*_0_ is the average time it take a protein to diffuse the length of the domain^42^. If the bounded domain diffuses normally, the equation can be modified so that,

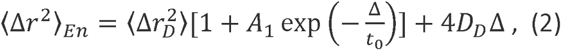

where *D*_*D*_ is the diffusion constant of the domain. Fitting Eq. 2 to **Figure 4C** we obtain that 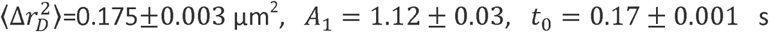, *A*_1_ *=* 1.12 ±0.03, *t*_0_ = 0.17 ± 0.001 s, and *D*_*D*_ *=* 0.01 ± 0.001 μm^2^/s. Interestingly, similar domain areas were found for different cell types ^42^, for example, a study using super-resolution imaging to measure the distribution of area size of the cortical actin networks found a wide distribution of area sizes peaked at 0.1-0.2 μm^2^ ^6^-Assuming proteins undergo normal diffusion within the confined domain we can extract the diffusion coefficient of TrkB and p75 to be 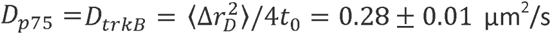, which is 28 times larger than the diffusion constant of the confining domain. Moreover, we observe a second change in the MSD slope starting at Δ∼2.5 s, which gives an estimate for the residency time in the confinement domain. The time and ensemble averaged MSD, ⟨ Δ*r*^*2*^ ⟩_*TE*_, of both receptors is shown in Figure **4D** The proteins exhibit subdiffusion, ⟨Δ*r*^2^⟩_*TE*_ *∼* Δ^α^ with a power law α *∼* 0.73 for A > 0.05 s. We note that the time and ensemble averaged MSD, ⟨ Δ*r*^2^ ⟩_*TE*_, is qualitatively different from ⟨ Δ*r*^2^ ⟩_*En*_ (Figure **4D),** indicating a non-ergodic diffusion mechanism ^33,43^. Similar subdiffusion behavior with similar power laws was observed previously, for example for ion channels in Human embryonic kidney (HEK) cells^6,32^.

**Figure 4:**
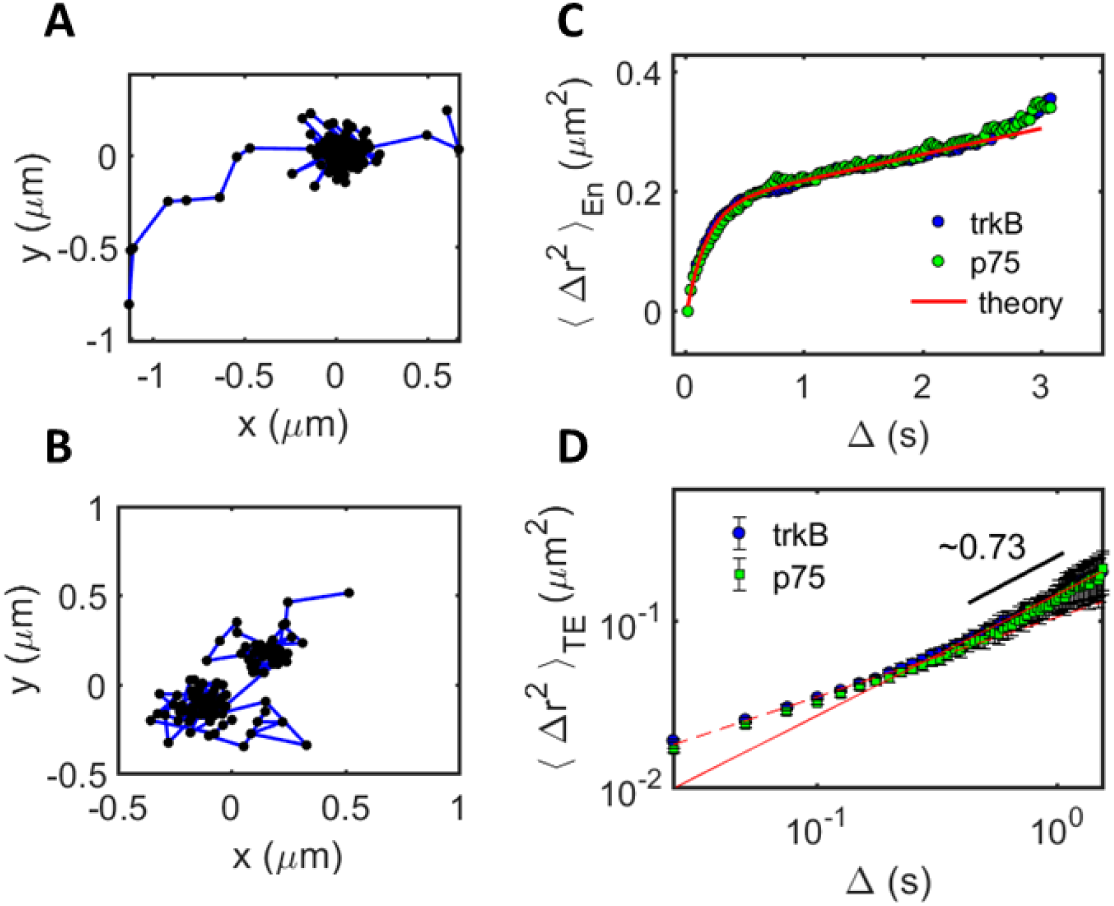
Transient confinement in the motion of TrkB receptors in the plasma membrane of neuronal cells. Typical trajectory of **A** TrkB receptor **B** p75 receptor. **C** Mean square displacement of TrkB (green triangles) and p75 (purple circles) showing confinement in sub-micron domains for durations of several seconds.

To further support the conclusion that TrkB and p75 are confined to physical domains within the membrane, we plot the probability distribution of finding a receptor at any location within the membrane during the entire span of an experiment, as shown in Figure 5A. Similar domains were found when studying the spatial distribution of dendritic cell-specific intercellular adhesion molecule-3-grabbing nonintegrin in Chinese hamster ovary cells^47^. The analysis shows clear domains in which the occupation probability is high. A closer examination of the trajectories contributing to each domain, as shown in Figure 5B for the highlighted domain in Figure 5A reveals that receptors entered the same domain at different times, the first entering 0.4 s after the recording started and the last receptor entered 11 s after the recording started. We were able to observe in several cases two receptors entering the same domain, or a receptor entering an occupied domain. However, our tracking algorithm is unable to follow the motion of two proteins in the same domain, since their fluorescence signal overlaps most of the time in such a small domain. We conclude that the observed domains are actual locations within the membrane, and that their lifetime is at least as long as our experiment length, which is 15 s.

**Figure 5:**
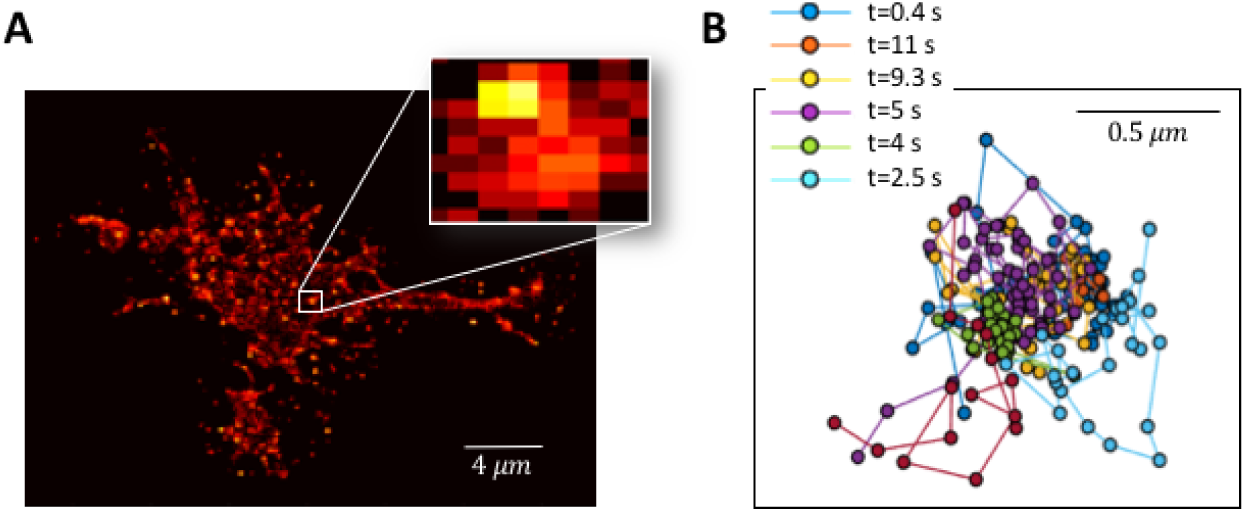
Characterization of compartments in the plasma membrane from the trajectories ofp75. A The probability distribution of finding a receptor at any time during the experiment. Domains of higher probability are present throughout the cell, one of each is shown at higher magnification. B trajectories of different receptor entering the domain in the inset of A. Each receptor enters the domain at a different time as indicated in the figure.

## Discussion

The emerging picture from our combined results is that flows in the plasma membrane induced, for example, by the motion of transmembrane proteins, propagate through the membrane at intermediate distances but are arrested on a length scale equivalent to the size of these domains. Such a severe suppression of flow can result only from momentum absorption due to an immobile inclusion around which the plasma lipids cannot flow. We note that a few immobile obstacles cannot account for this flow suppression, they will affect the amplitude but not the long range power low decay of the flow field^37^.

We also find that there are physical domains in the plasma membrane. Protein receptors can diffuse into these domains and escape from them. In addition, these domains maintain their integrity for, at least, several seconds. The domains themselves diffuse at a low rate. It is known that domains in the plasma membrane can originate from various sources, such as lipid microdomains sensitive to cholesterol^35^ and sphingomyelin concentration ^48^ or interactions with the underlying actin cytoskeleton (pickets and fences model)^8^’^48^. In addition, the motion of proteins can be affected by interactions with other biomolecules ^10^·^47^·^49^. Most probably, the motion of plasma proteins is susceptible to all these effects. It is yet unclear whether the confining domains observed here are the cause of the suppression of flow in the plasma membrane. Experiments to elucidate the source of flow suppression are currently underway.

One possible interpretation of these results is that the domain boundaries are surrounded by relatively immobile obstacles (e.g. proteins anchored to underlying actin cortical mesh) which form dense enough fences to suppress lipid flow (see Figure 6). Such an interpretation is consisted with other studies ^6^. Since the actin network fluctuates and changes as well as the internal section of the receptors, it is expected that these domains will diffuse and change in time on a longer timescale. This interpretation is supported by the high protein density in the plasma membrane and the many reports that implicate that the cortical actin network plays a role in restricting protein motion in the cell membrane (for example ^6,48,50-55^). Moreover, studies of the structure of the actin cortex^6,56^ show that the actin mesh size in comparable to the length scale emerging form our various measurement for domain size and flow suppression, i.e. ∼ 0.4 *μm.*

**Figure 6:**
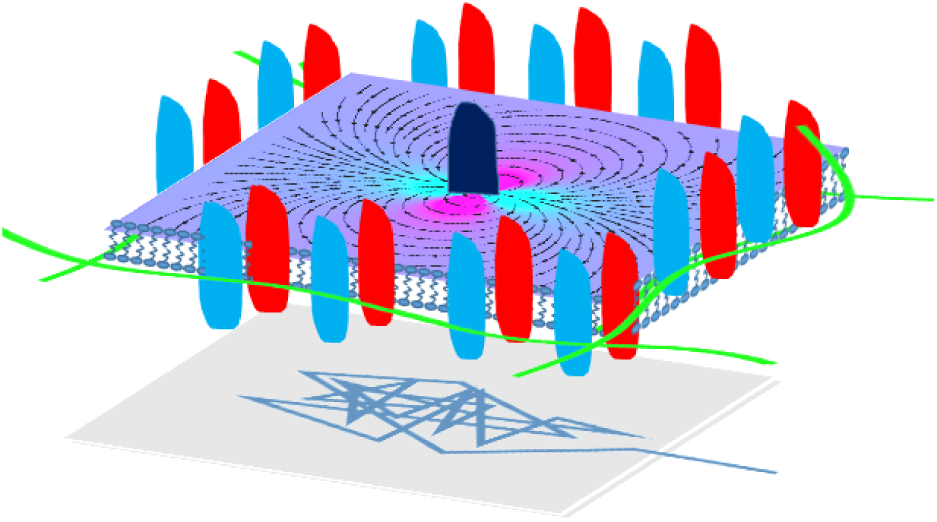
An optional model for flow arrest due to dense protein arrangement entangles with the underlying actin mesh. The flow field in the plasma membrane induced by protein motion decays strongly due to momentum absorption by immobile proteins. The trajectory of the tracer protein (depicted below the membrane) reflects the transient local trapping of the tracer protein.

Our results and suggested interpretation provide a microscopic explanation to the surprising finding that local changes in the plasma membrane result only in a local perturbation of membrane tention^21^. Combined, the observation that perturbations to membrane tension do not propagate throughout the cell have significant implication to models relying on mechanical signaling across cells.

From the vast literature concerned with the plasma membrane structure it is clear that the factors governing its local organizations are multiple, and that their combined effect is complex. How these factors affect and contribute to optimize sensing and signaling at the interface between a cell and its surrounding is still not fully understood. However, some studies hint that the passive receptor confinement in microcompartments of the plasma membrane may promote the maintenance of signaling protein complexes^55^. We hypothesize that the correlated motion of proteins within these compartments may add to the integrity of such complexes without affecting proteins at larger distances.

## ACKNOWLEDGMENTS

We thank Haim Diamant for helpful discussions. This work was supported by the ISF (Israel Science foundation) grants 561/11 and 988/17, and the ERC (European Research Council) grant 309377. We thank Eitan Erez Zahavi and Tal Gradus-Perry for technical help with TIRF imaging and mice colony.

## AUTHOR CONTRIBUTIONS

M.C. performed the experiments and part of the analysis. E.P. designed and supervised the experiments. Y.R. conceives the project, lead the analysis and wrote the manuscript.

## Methods

All methods were carried out in accordance with Tel-Aviv University guidelines and regulations. All animal experimentations were approved by Tel-Aviv University Animal Ethics Committee.

### Dissociated spinal cord culture

Neuronal cell culture was performed according to “Rapid purification of embryonic rat motor neurons: an in vitro model for studying MND/ALS pathogenesis” (1). Briefly, spinal cords were collected from C57BL/6 or ICR mice at E12.5. In all experiments, mice of both sexes were used. Spinal cords were then dissociated using trypsin and repeated trituration. Cells were collected through centrifugation followed by Optiprep (Sigma-Aldrich) gradient centrifugation to achieve a motor neuron-enriched cell culture. Neurons were then plated on glass bottom dishes (10,000 cells per insert) coated with complete Neurobasal medium as describe before^57^. All neuronal cultures were grown for 4 DIV.

### Preparation of glass bottom culture plate samples

Cells were plated on round coverslip-bottom 35mm dishes with #1.5 thickness (FD35-100, WPI). Before use, the plates were cleaned as follows: Plates were treated with 20% NaOH for 45’, washed in DDW, then treated with 30% sulfuric acid for 30’, rinsed in DDW and cleaned with 70% ethanol then washed with sterile DDW and left to dry in a sterile hood. Before plating cells, PDMS was mixed and cast in round plates, then cut to fit on the glass bottom dishes. Wells for plating the cells were punched using 6mm (4 wells / plate) puncher. PDMS wells cast was cleaned with adhesive tape and 70% ethanol, and left to dry inside a biological hood. The cast was firmly attached to the glass surface and the plate was incubated for 5-10’ in 60°C. PDMS was pressed against the glass to ensure tight bondage. For plating spinal cord neurons, wells were treated over night with poly-ornithine and 2 hours of laminin. For plating HEK293T cells, wells were treated with 0.1% poly-L-lysine (P4707, Sigma Aldrich) for 45’ before plating. HEK293T cells were maintained in DMEM (Biological Industries, Israel) supplemented with 1% Glutamax (Gibco), 10% Fetal Bovine Serum and 1% Penicilin-Streptomycin (Biological Industries, Israel). Neurons, were plated at a density of 10,000 cells/6mm PDMS well. HEK293T were plated on 8mm wells at a density of 15,000/well.

### Preparation of expression plasmids for the study of membrane receptors

TrkB-GFP plasmid, encoding rat full-length TrkB fused to EGFP at the C’-terminus under a CMV promoter was gifted by Rosalind Segal (Harvard University). p75-GFP plasmid, encoding rat p75NTR fused to EGFP was a gift by <u>Francisca C Bronfman</u> (<u>Pontifical Catholic University of Chile</u>). The pLL3.7-CMV-EGFP 3rd generation lentiviral vector (Addgene #11795) was gifted by Uri Ashery (Tel Aviv University). LV-TrkB-GFP and LV-p75-GFP were cloned by inserting TrkB and p75 from TrkB-GFP and p75-GFP into pLL3.7-CMV-EGFP downstream of the CMV promoter. LV-TrkB-ACP was subcloned by inserting the ACP sequence from pACP-tagm2 (NEB) immediately after the TrkB signal sequence and cloning into pLL-3.7-CMV whose EGFP reporter was excised.

### Plasmid expression in HEK293T and spinal cord neurons

Lentiviral particles were produced in HEK cells using a 2^nd^ generation packaging system based on Gag-Pol helper and VSVG coat constructs with selected Lentiviral (LV) vector construct. For lentiviral production, HEK cells grown on 60mm dish at 70-80% confluence were transfected 10μg of LV-vector together with 7.5μg of pGag-Pol and 2.5μg of pVSVG. Plasmids were mixed in Calcium-Phosphate transfection mix: 25mM Hepes, 5mM Kcl, 140mM NaCl, 0.75mM Na_2_Po_4_ with 125mM CaCl_2_ immediately before addition to cells, in a volume of 0.5mL per plate. Culture supernatants were harvested at 2 days post-transfection, and concentrated ×10 using the PEG virus precipitation kit (Abeam, abl02538). Final pellet was re-suspended in Neurobasal media, aliquoted and kept in −80°C until use. For SCN transduction, 2-10μl of concentrated LV suspension was used per well containing 10,000 neurons. LV were added 1-2 hours after plating of the neurons, and were washed out three times in CNB 24 hours later. For HEK live and fixed cell imaging experiments, cells were plated and transfected 1 day post plating with Fugene 6 reagent in DMEM.

### Total Internal Reflection Fluorescence (TIRF) and Oblique fluorescence microscopy

Live and fixed cell TIRF imaging was done on a FEI-Munich iMic-42 digital microscope equipped with fast 360° spinning beam scanner to allow even illumination of the entire diameter of the back focal plane of the objective. A 100 × Olympus 1.49 numerical aperture TIRF objective was used for objective-based TIR. As illumination source, 4 solid-state laser lines at 405, 488, 561 and 640nm were used with maximum output power of 50mW each. Control of stage, excitation and acquisition parameters were via Live Acquisition 2 software. Images were captured using Ixon897 EMCCD camera (Andor). In all live imaging experiments, a 37°C, 5% C02 and humidity conditions were kept using a custom environmental control system (Live Imaging Services), and gain was set to 300 to maximize signal capture and minimize exposure of the sample. For SPT experiments, exposure time was 25 milliseconds with 1 millisecond delay. Imaging area was cropped prior to imaging to allow fast imaging times. Laser intensities used were 40% and 70% for 561 and 488 respectively. TIRF angle varied for each plate but was between 2.480-2.500. all movies acquired were 1500 frames long.

### Surface ACP-CoA labeling

Freshly prepared mixture of ACP-CoA labeling, New England Biolabs (NEB), was used for single-dye ACP labeling. 2μM fluorescent CoA (488, 547 or 647), ImM MgCI2, 0.8nM SFP-synthase enzyme in either DMEM with 1% Glutamax (for HEK cells) or Neurobasal 2% B27, 1% Glutamax (for spinal cord neurons) supplemented with 0.5% BSA. A volume of 25 μl was added for 6 mm PDMS wells, and incubated 30’ in the cell-culture incubator. Cells were then washed 3 times with the media.

### Mean square displacement analysis

Single particle trajectories were extracted from the TIRF movies by conventional video microscopy (2). For MSD analysis we used all mobile particle trajectories, defined so that their maximum displacement from their original position was larger than 0.5 *μm.*

The ensemble average mean squared displacement of tracer particles is defined as the average over all particles of the squared displacement of each particle at time τ from its position at τ = 0:

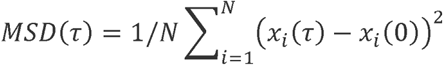

where *x*_*i*_(τ) is time the position of particle *i* at time τ, and *N* is the total number of tracked particles.

The ensemble and time average mean squared displacement of tracer particles is defined as the average over all particles of the squared displacement of each particle at time τ from its position at τ = 0:

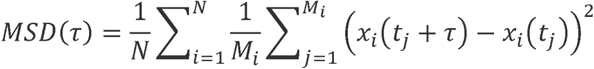

where t_j_ is the acquisition time series, τ is the lag time (the time over which the squared displacement is measured), and *M*_*i*_ is the total number of trajectory segments of length τ extracted for particle *i.* To obtain the plots in Figure 4C,D we averaged particles in different locations within the cell membrane, in different cells, and for different cultures.

