## Supplemental Materials for "Flow arrest in the plasma membrane"

**Supplementary material**

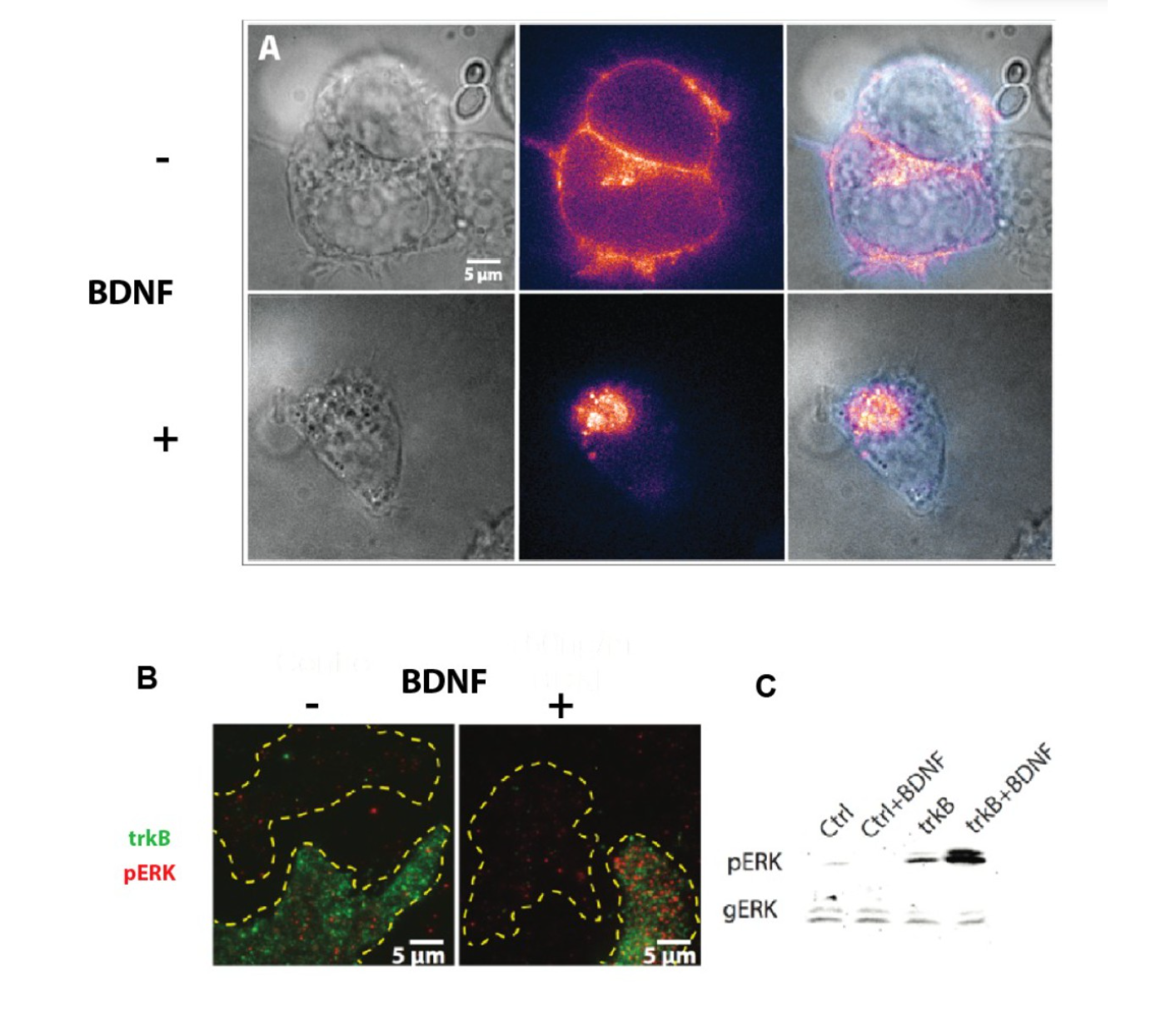

***Supplementary figure 1: TrkB ACP is biologically active. A)*** *TrkB ACP expressing motor neurons were imaged 30 minutes after the addition of 50 ng/ml BDNF or control media. Results indicate that TrkB-ACP is internalized in response to 50 ng/ml BDNF within 30 minutes. Image was acquired using pseudoTIRF (Cui et al. 2007).* ***B)*** *TIRF images showing HEK cells that express TrkB ACP (green) respond to 50 ng/ml BDNF treatment by activation of pERK (red) more strongly then non ACP expressing cells (non-green cells marked by yellow outline). Cells were fixed 30 minutes after treatment, permibilized using triton and stained with Sigma's mouse anti pERK antibody (M8159) at 1:5000 concentration.* ***C)*** *Immunoblots demonstrating the activation of pERK in HEK cells with/out TrkB ACP transfection and with/out 50 ng/ml BDNF treatment. Cells were lysed 30 min after treatment, and prepared for blotting. Antibodies used were Sigma's mouse anti pERK antibody (M8159) at 1:5000 concentration, and Sigma's rabbit anti general ERK antibody (M5670) at 1:10000 concentration.*

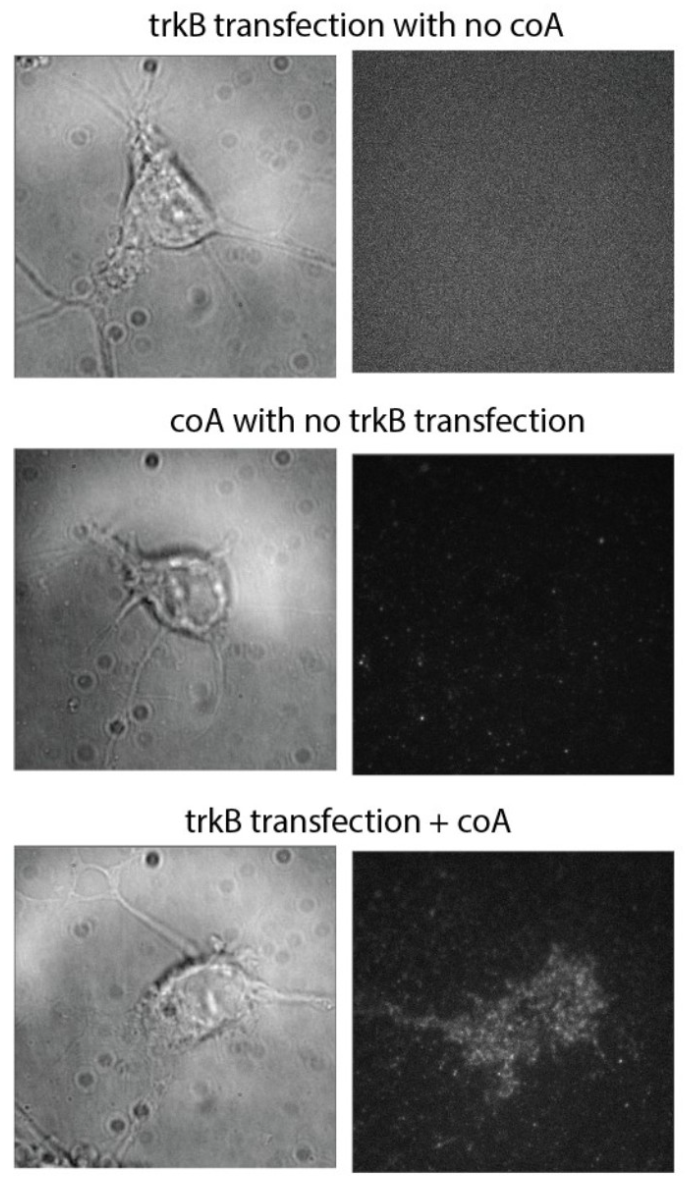

***Supplementary figure 2: The specificity of the ACP tag*** *– Left panel shows bright field images of motor neuron, while he right panel shows the same field of view in TIRF, with the 488 laser set to 70%. As shown, the background noise is very minute, and only the condition in which cells were transfected and the dye was added, show specific signal.*
